# Stressor-specific dynamic patterns of noradrenaline release in the paraventricular nucleus of the hypothalamus in freely moving mice

**DOI:** 10.64898/2026.07.25.740743

**Authors:** Ryunosuke Ono, Keiichi Itoi, Kenji Sakimura, Takuya Sasaki, Hiroyuki Igarashi

## Abstract

An organism is constantly challenged with various stressors. These stress signals ultimately converge on the hypothalamic paraventricular nucleus (PVN), where they are integrated by corticotropin-releasing hormone (CRH)-producing neurons that are primarily involved in the regulation of the hypothalamic-pituitary-adrenal (HPA) axis. Noradrenaline (NA), among others, is recognized as the major transmitter that regulates the PVN-CRH neurons, and thereby is involved in the regulation of the HPA axis as well as the autonomic outflow. Previous studies have demonstrated that stress increases NA release within the PVN^1^ and that NA activates CRH neurons^2^. However, NA release patterns from the axon terminals in the PVN upon stress exposure have not yet been studied, because continuous monitoring of transmitter release became possible only recently.

In the present study, we aimed to monitor stressor-dependent NA release patterns in the PVN. To continuously monitor NA release in freely moving mice, a fluorescent NA sensor (GRAB_NA_)^3,4^ was expressed in the PVN, and the emitted signals were recorded via fibre photometry. Following stress exposure, NA release took place over distinct timescales ranging from seconds to hours. For example, a pulsatile NA release was observed in seconds in response to an acute nociceptive stressor. In contrast, sustained physical stressors, such as tail suspension and restraint, induced much more prolonged NA release that persists throughout the stress exposure period over tens of minutes. Intraperitoneal administration of lipopolysaccharide produced a gradual increase in NA release that persisted for several hours. These findings demonstrate that NA is released into the PVN in a stress-specific manner. In addition, elevated NA release was associated with increased behavioural activity, characteristic of each stressor. Together, these findings provide a framework for understanding how the temporal dynamics of NA release in the PVN encode diverse stress signals, and regulate neuroendocrine outputs.

## 1. Introduction

Living organisms are constantly exposed to a wide variety of stressors arising from both internal and external sources. These stressors elicit stress responses that facilitate adaptation and survival, and maintain internal homeostasis^5^. Stress responses occur across multiple timescales: a rapid increase in autonomic output induces pupillary dilation and alters cardiovascular dynamics to increase blood and oxygen supply to skeletal muscles and other vital organs, accompanied by heightened arousal behaviours. The neuroendocrine stress response promotes gluconeogenesis and glycogenolysis, as well as transient immune modulation mediated by glucocorticoids. Collectively, these coordinated physiological changes constitute the stress response, which is essential for survival and adaptation.

Stress-induced neuronal signals originate in multiple brain regions and ultimately converge on the paraventricular nucleus of the hypothalamus (PVN). Corticotropin-releasing hormone (CRH)-producing neurons (hereafter CRH neurons) in the PVN receive stress-responsive inputs, and in turn, regulate both the autonomic nervous system and the hypothalamic–pituitary–adrenal (HPA) axis. Excessive release of CRH leads to chronic hypersecretion of glucocorticoids^6^ and has been implicated in stress-related psychiatric disorders, including post-traumatic stress disorder (PTSD)^7^ and depression^8^, as well as metabolic disorders such as diabetes and hypertension. Thus, elucidating the mechanisms regulating CRH neuron activity *in vivo* is a critical step towards understanding stress-related pathophysiology and developing pharmacological strategies for the treatment of stress-related disorders.

Noradrenaline (NA) has long been recognized as a key neurotransmitter regulating CRH neuronal activity^9^. NA is released from brainstem nuclei such as the locus coeruleus (LC) and the nucleus of the solitary tract (NTS), which project broadly to brain regions including the hypothalamus, the cerebral cortex, and the hippocampus^10,11^. Stress exposure has been shown to trigger NA release within the PVN^1^, and studies using acute brain slices have demonstrated that NA activates CRH neurons^2^, although it is controversial whether NA action is directly on CRH neurons^12^ or is mediated by hypothalamic interneurons^13^. While these findings strongly suggest that NA enhances CRH neuronal activity, dynamic patterns of NA release *in vivo* remain poorly understood. Given the considerable variability of stressors in terms of modality and duration, stressor-dependent modulation of NA release may be necessary for appropriate stress responses, which are partly mediated by CRH neuronal activity. However, conventional methods such as microdialysis lack both the temporal and spatial resolution necessary for monitoring NA release within a discrete brain region such as the PVN; more specifically, microdialysis requires several minutes to collect extracellular fluid samples, making it unsuitable for monitoring rapid fluctuations occurring in the order of seconds^14^, and its spatial resolution of approximately 1 mm is insufficient to monitor NA release within a smaller-brain region such as the PVN. These technical limitations underscore the need for a novel approach capable of continuously monitoring localized NA release *in vivo* with high temporal and spatial resolution.

To address these limitations, we established a recording system combining a GPCR-activation-based (GRAB) fluorescent sensor with fibre photometry to monitor NA release *in vivo*. The GRAB_NA_ sensor is based on a modified α2-adrenergic receptor into which a circularly permuted enhanced green fluorescent protein (cpEGFP) has been inserted within the third intracellular loop^3,4^. Upon binding of NA to the ligand-binding domain, a conformational change is induced in the sensor, resulting in an increase in cpEGFP fluorescence intensity. By genetically expressing the GRAB_NA_ sensor in specific cell types, this approach enables real-time monitoring of NA release at the site of expression, measured as changes in observed fluorescence intensity. This approach provides improved temporal and spatial resolution, enabling detection of fluctuations of NA release in the order of hundreds of milliseconds with micrometre-scale spatial precision^15^.

In the present study, we employed this recording system to monitor NA release in real time in mice exposed to a variety of stressors with distinct temporal characteristics, to examine whether NA release exhibits stressor-specific temporal patterns. Our findings reveal that NA is released into the PVN in a stressor-dependent manner, providing new insights into the temporal coding of stress information in the hypothalamus.

## 2. Methods

### Approvals

All experiments were performed with the approval of the Animal Experimentation Ethics Committee at Tohoku University. (approval number: 2022 PhA-004) and in accordance with the NIH guidelines for the care and use of animals.

### Animals

C57BL/6N mice (eight weeks old) were used. All wild-type mice were obtained from SLC (Shizuoka, Japan). CRF-iCreΔNeo mice^16–20^ were generated by K.S. and K.I. (Supplementary Figure 1). Mice were housed under controlled conditions (23 ± 1 °C; 50 ± 5% relative humidity) with a 12-h light/12-h dark cycle (lights off at 8:00 PM) and had ad libitum access to food and water. They were maintained with free access to water and food under a 12-h light/12-h dark schedule (lights off at 8:00 PM), with housing conditions at 23 ± 1 °C and a relative humidity of 50 ± 10%

### Reagents

Lipopolysaccharide (LPS; Escherichia coli K-235, Sigma-Aldrich, USA) was administered intraperitoneally at a dose of 0.5 mg/kg. Yohimbine hydrochloride (Y3125-1G; Sigma-Aldrich, USA) was administered intraperitoneally at a dose of 2.0 mg/kg.

### Surgery

Mice were anesthetized with 3% isoflurane and maintained with 1–3% isoflurane throughout surgery. After confirming suppression of reflexes, the head was secured in a stereotaxic apparatus (SMM-200; Narishige, Tokyo, Japan). Veterinary ointment (Propeto; Maruishi, Osaka, Japan) was applied to the eyes to prevent corneal drying. Prior to each incision, the skin was sterilized with betadine followed by 70% ethanol. The scalp was incised along the midline using surgical scissors to expose the skull, and connective tissue was removed while applying 2% hydrogen peroxide to the skull surface. A small craniotomy was made at the target coordinates using an electric drill fitted with a 0.5-mm diameter round drill bit.

For virus injection, a glass pipette (f = 10 μm) mounted on a syringe pump (Legato 111; Muromachi Kikai, Tokyo, Japan) was used to deliver viral vectors into the target brain regions. Viruses were infused at a rate of 100 nL/min, and the pipette was left in place for 10 min after injection before being slowly withdrawn. Stereotaxic coordinates were as follows: NTS (mm from obex; AP, −0.5; ML, ±0.2; DV, −0.5), LC (mm from bregma; AP, −5.5; ML, ±0.8; DV, −2.75), and PVN (mm from bregma; AP, −0.7; ML, −0.2; DV, −4.7).

Injection volumes were 100 nL/site for AAV2-retro-CaMKII-Cre, AAV8-hSyn-DIO-mCherry, and AAV1-hSyn-FLEX-EGFP, and 300 nL/site for AAV9-hSyn-GRABNA2h/2m (PT-5262, PT-1344; BrainVTA, China) and AAV9-hSyn-GRABNAmut (PT-1343; BrainVTA, China).

For optical fibre implantation, the skull surface was thinned using a bone scraper (10075-16; Fine Science Tools, Germany), and four stainless-steel screws (diameter, 1.0 mm; length, 2.0 mm) were placed as anchor screws. Following viral injection, an optical fibre (400 μm diameter; R-FOC-BF400c-39NA, RWD, China) was inserted at a rate of 1 mm/min and targeted to the PVN (mm from bregma; AP, −0.7; ML, −0.2; DV, −4.5). The optical fibre was secured to the skull using UV-curable adhesive (B-7202P; BISCO, USA), which was polymerized by ultraviolet illumination. Dental cement was then applied over the implant for additional stabilization.

Following surgery, anesthesia was discontinued and mice were allowed to recover on a heating pad with body temperature maintained. To minimize postoperative pain and distress, kanamycin (100 mg/kg) was administered subcutaneously, and meloxicam (5 mg/kg) was administered intraperitoneally together with saline adjusted to compensate for body weight loss. Following surgery, each animal was housed individually in a transparent Plexiglas cage with free access to food and water.

### Immunohistochemistry

The mice were overdosed with isoflurane, transcardially perfused with 4% PFA in PBS, and decapitated. After dissection, the brains were fixed overnight in 4% PFA and then equilibrated in 30% sucrose in PBS for an additional night. Frozen coronal sections (50 μm) were cut using a cryostat (CM1950; Leica, Wetzlar, Germany) and mounted onto glass slides (S9901; Matsunami Glass Ind., Japan). Brain slices were washed three times in PBS for 5 min each. Slices were permeabilized in PBS containing 0.3% Triton X-100 and 5% bovine serum albumin (BSA) at room temperature for 60 min.

For EGFP and mCherry immunostaining, slices were then incubated overnight at 4 °C with the following primary antibodies: chicken anti-GFP (1:1000; Abcam, ab13970, UK), and rabbit anti-DsRed (1:1000; Takara Bio, 632496, USA). After rinsing with PBS, slices were incubated for 90 min at room temperature with Alexa Fluor 488-conjugated anti-chicken IgG (1:1000), and Alexa Fluor 594-conjugated anti-rabbit IgG (1:1000) (Jackson ImmunoResearch, USA) in PBS containing 0.3% Triton X-100. Sections were then stained with DAPI (1:1000; D9542-1MG; Sigma-Aldrich, USA) for 15 min, followed by three additional PBS washes (5 min each). Samples were mounted with Fluoro-KEEPER Antifade Reagent (Nacalai Tesque). Images were acquired using a fluorescence microscope (BZ-X1000; Keyence, Osaka, Japan) equipped with a water-immersion objective lens (×10, 0.75 NA). For dopamine β-hydroxylase (DBH) immunostaining, anti-DBH antibody (1:500; JH3126; provided by K.I.) was used as the primary antibody, and Alexa Fluor 594 anti-rabbit IgG (1:500; 711-586-152; Jackson Laboratory, USA) was used as the secondary antibody.

### Fibre photometry

A fibre-coupled blue LED with an integrated driver (470 nm; M470F4; Thorlabs) was used as the light source for the photometry system. The excitation light intensity for GRAB_NA_ was set to 25 µW at the fibre tip. The 470 nm excitation light passed through a bandpass filter (86-341, 466 ± 20 nm; Edmund Optics) and was reflected by an excitation dichroic mirror (67-064, 506 nm; Edmund Optics) before being delivered through the fibre. The emitted green fluorescence signal was filtered through a dichroic mirror (67-066, 562 nm; Edmund Optics) and a bandpass filter (67-016, 520 ± 18 nm; Edmund Optics), restricting detection to 513–538 nm. The signal was then detected using a photomultiplier tube (PMT; CR131; Beijing Hamamatsu Photon Techniques Inc.). Signals from each PMT were digitized using a data acquisition board (COME2-FTR-PYP; Lucir) and recorded on a PC using pyPhotometry^21^ for acquisition settings and data logging. Photometry data were sampled at 100 Hz and analyzed using custom Python scripts. First, noise was reduced using a smoothing filter. Next, a regression model was fitted to each signal to correct for photobleaching, and the slope was adjusted accordingly. The fluorescence change (ΔF/F) was then calculated using the mean signal during the first 30 s after the start of recording as the baseline. These processed data were subsequently used to compute the relevant parameters.

### Foot shock

Foot shock stimulation was conducted in a custom-made acrylic chamber (215 mm × 215 mm × 300 mm). After connecting the mouse to the optical fibre for photometry recording, the animal was transferred from its home cage to the stimulation chamber, where stimulation was delivered under freely moving conditions.

Electric shocks were delivered at 0.8 mA using a shock generator (SGS-003DX; Muromachi Kikai, Tokyo, Japan). A cycle timer (CBX-CT; Muromachi Kikai, Tokyo, Japan) was used to deliver 2-s shocks at 3-min intervals. After completion of the recording, the mouse was removed from the chamber and returned to its home cage. Prior to the recording sessions, mice were habituated to the stimulation chamber by allowing free exploration for 15 min per day for three consecutive days.

### Tail pinch

Nociceptive stimulation was applied by pinching the tail of the mouse in its home cage for 2 s using forceps (SP 133; Sonne, Tokyo, Japan). The interstimulus interval was set to 3 min, consistent with the foot shock protocol. The duration and timing of each stimulus were manually recorded using a stopwatch synchronized with the photometry recording.

### Tail suspension

Mice were removed from their home cage and suspended by the tail for either 1 min or 10 min. For the 1-min condition, the distal end of the tail was held manually by the experimenter to keep the mouse immobile. For the 10-min condition, a custom plastic tube (1.5 cm in diameter, 4 cm in length) was placed over the tail^22^, and the distal end of the tail was secured with tape to a stainless-steel bar (1 cm in diameter, 56 cm in length) positioned 50 cm above the floor. Behaviour was simultaneously recorded using a video camera (HDR-CX680; Sony, Tokyo, Japan). The recorded videos were downsampled from 30 frames per second (fps) to 1 fps, and periods during which no movement was detected between consecutive frames were quantified as immobility time.

### Restraint

Mice were removed from their home cage and placed into a cylindrical restrainer (KN-325-C-1; Natsume Seisakusho, Tokyo, Japan). The restrainer containing the mouse was then returned to the home cage, and the mice were restrained for 10 min.

### Open-field test

Mice were removed from their home cage and placed in a black cubic open-field arena (42.5 cm per side), where they were allowed to freely explore for 10 min. Behaviour was recorded using a video camera (HDR-CX680; Sony, Tokyo, Japan).

The recorded videos were downsampled from 30 frames per second (fps) to 1 fps, and the trajectory and total distance travelled were analyzed using the Manual Tracking function in ImageJ. The floor of the open-field arena was divided into nine equal sections (3 × 3 grid), and the time spent in the central section was defined as the centre time.

### Data analysis

Statistical analyses were performed using GraphPad Prism 10 (GraphPad Software, USA). All data are presented as mean ± standard error of the mean (SEM). Statistical comparisons were conducted using Student’s *t*-test, paired *t*-test, one-way ANOVA, or Friedman’s test, as appropriate. Statistical significance was defined as *p* < 0.05.

## 3. Results

### PVN receives predominant projections from the NTS

The PVN receives neural inputs from multiple brain regions. Among these, we anatomically characterised the major nuclei sending noradrenergic projections to the PVN. A previous study using a neuron-specific anterograde tracer demonstrated that the PVN receives noradrenergic projections from medullary regions, including the nucleus of the solitary tract (NTS), as well as from the locus coeruleus (LC)^10^. In the present study, we selectively visualized upstream projection pathways to the PVN using a two-step viral vector strategy.

First, an adeno-associated virus (AAV) vector expressing the recombinase Cre in a retrograde manner (retroAAV-Cre)^23^ was injected into the PVN of mice. One week later, a Cre-dependent AAV vector expressing enhanced green fluorescent protein (EGFP) was injected into the NTS, while a Cre-dependent AAV vector expressing mCherry was injected into the LC (Figure 1A). This strategy enabled selective labelling of NTS neurons projecting to the PVN with EGFP, and LC neurons projecting to the PVN with mCherry.

**Figure 1:**
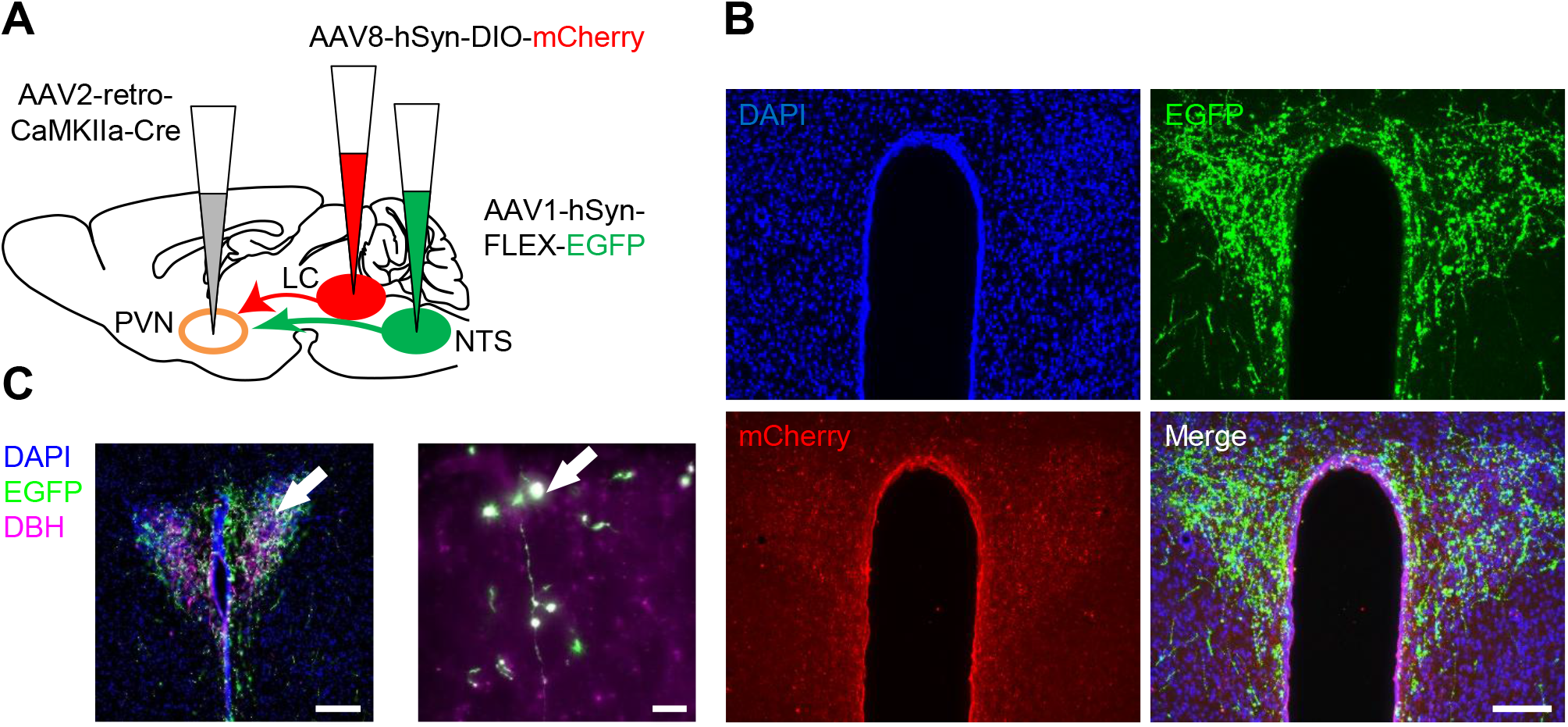
PVN receives predominant projections from the NTS. A. Schematic illustration of experimental design. B. Representative images of DAPI, EGFP, and mCherry staining in the PVN. Scale bar = 100 μm. C. Representative images of DAPI, EGFP, and DBH staining in the PVN. The right panel shows a magnified view of the area indicated by the arrow. Left panel: Scale bar = 100 μm; right panel: Scale bar = 10 μm.

Immunohistochemical analysis of the PVN revealed robust EGFP-positive fibres within the nucleus, whereas mCherry-positive fibres were scarcely detected (Figure 1B). To further confirm whether the NTS-to-PVN projection was noradrenergic, we performed immunostaining for dopamine β-hydroxylase (DBH), a rate-limiting enzyme in noradrenaline biosynthesis. Colocalization of EGFP-positive fibres with DBH immunoreactivity was observed within the PVN (Figure 1C).

Taken together, these results indicate that noradrenergic projections to the PVN predominantly originate from the NTS rather than the LC.

### Transient nociceptive stress induces NA release on a second timescale

Next, we examined how NA release in the PVN changes in response to stress exposure. To detect NA release, we expressed a GPCR Activation-Based fluorescent sensor for noradrenaline (GRAB_NA_)^3,4^ in neurons of the PVN. Following the injection of an AAV vector expressing GRAB_NA_ into the PVN of mice, an optical fibre was implanted directly above the PVN (Figure 2A). Fluorescence changes from the GRAB_NA_ were recorded using the fibre photometry system in freely moving animals (Figure 2B, C).

**Figure 2:**
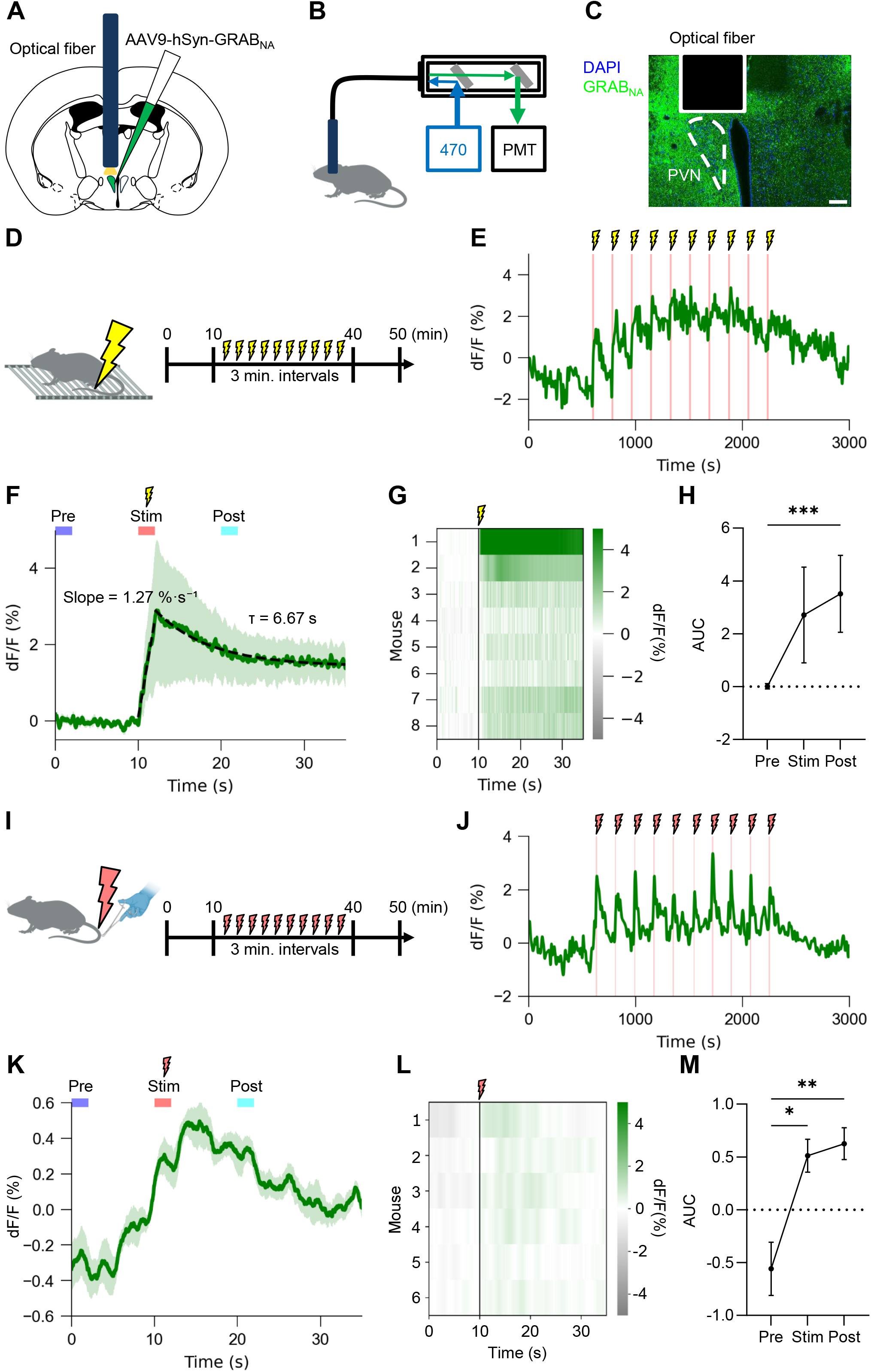
Transient nociceptive stress induces NA release on a second timescale. A. Schematic illustration of virus injection and fibre implantation to the PVN. B. Schematic diagram of the fibre photometry recording. C. Representative image showing the expression of GRAB_NA_ and the fibre tract. Scale bar = 100 μm. D. Schematic illustration of the foot shock stimulation paradigm. E. Representative trace showing NA release throughout the foot shock recording session. Red lines indicate the timing of foot shock stimulation. F. Averaged traces showing NA release before and after foot shock stimulation. (Male, *N* = 4; female, *N* = 4; 10 trials per animal; mean ± SEM) G. Heat map showing individual NA release before and after foot shock stimulation. (10 trials per animal) H. AUC values for the Pre, Stim, and Post time bins. (Male, *N* = 4; female, *N* = 4; mean ± SEM; Friedman’s test; \*\*\**P* < 0.001) I. Schematic illustration of the tail pinch paradigm. J. Representative trace showing NA release throughout the tail pinch recording session. Red lines indicate the timing of tail pinch stimulation. K. Averaged traces showing NA release before and after tail pinch stimulation. (Male, *N* = 2; female, *N* = 4; 10 trials per animal; mean ± SEM) L. Heat map showing individual NA release before and after tail pinch stimulation. (10 trials per animal) M. AUC values for the Pre, Stim, and Post time bins. (Male, *N* = 2; female, *N* = 4; mean ± SEM; one-way ANOVA; \**P* < 0.05, \*\**P* < 0.01)

We first applied foot shock and tail pinch as acute nociceptive stressors. In the foot shock experiment, baseline activity was recorded for 10 min, followed by ten 2 s foot shocks (0.8 mA) delivered at 3 min intervals. Recording was continued for an additional 10 min following the final stimulation (Figure 2D). NA release increased in a time-locked manner with each foot shock stimulus, and the magnitude of NA release gradually accumulated with repeated stimulation (Figure 2E). To characterise the average response profile, traces representing the mean ± SEM of responses (averaged across the ten stimulations per animal) were plotted and subsequently averaged across animals (Figure 2F). NA release rapidly increased by approximately 3% immediately following stimulation onset, with the rising phase exhibiting a linear slope of 1.27 %·s⁻¹. The subsequent decay was well described by a single-exponential function with a time constant of 6.67 s, while the signal remained elevated by approximately 2% for the subsequent 20 s (Figure 2F). To visualize inter-individual variability, heat maps representing the averaged responses across the ten stimulations for each animal were generated (Figure 2G). Although all animals exhibited increased NA release following stimulation, the magnitude of the response varied across individuals (Figure 2G). For quantitative analysis, the area under the curve (AUC) was calculated for three periods: Pre (10 to 8 s before stimulation onset), Stim (the 2-s stimulation period), and Post (10 to 12 s after stimulation). The Post AUC was significantly greater than the Pre AUC (Figure 2H), consistent with stress-responsive NA release within the PVN.

In the tail pinch experiment, baseline activity was similarly recorded for 10 min, followed by ten 2 s tail pinches delivered at 3 min intervals using forceps applied to the tail of freely moving mice (Figure 2I). Similar to foot shock, NA release increased synchronously with each tail pinch stimulus (Figure 2J), with a preceding increase prior to stimulus onset (Figure 2K, L). Using the same definitions for Pre, Stim, and Post periods, AUC analysis revealed that both Stim and Post AUCs were significantly greater than the Pre AUC (Figure 2M).

These data demonstrate that both foot shock and tail pinch stimulation trigger rapid NA release on a timescale of seconds. Notably, while foot shock did not evoke preceding NA release, tail pinch induced an anticipatory gradual increase in NA release beginning approximately 10 s before stimulus onset.

### Sustained stress induces NA release on minute-to-hour timescales

We next investigated whether NA release patterns reflect the temporal characteristics of the applied stressors. After 10 min of baseline recording (Pre), animals were subjected to tail suspension for 10 min, defined as the Stim period. Recording continued for an additional 10 min following termination of tail suspension, defined as the Post period (Figure 3A). NA release progressively increased from the onset of tail suspension, reached a peak within several minutes, and remained elevated throughout the remainder of the stress period. Upon termination of tail suspension, NA release gradually declined toward baseline (Figure 3A, B). AUC quantification revealed that the Stim AUC was significantly greater than either the Pre and Post AUCs (Figure 3C).

**Figure 3:**
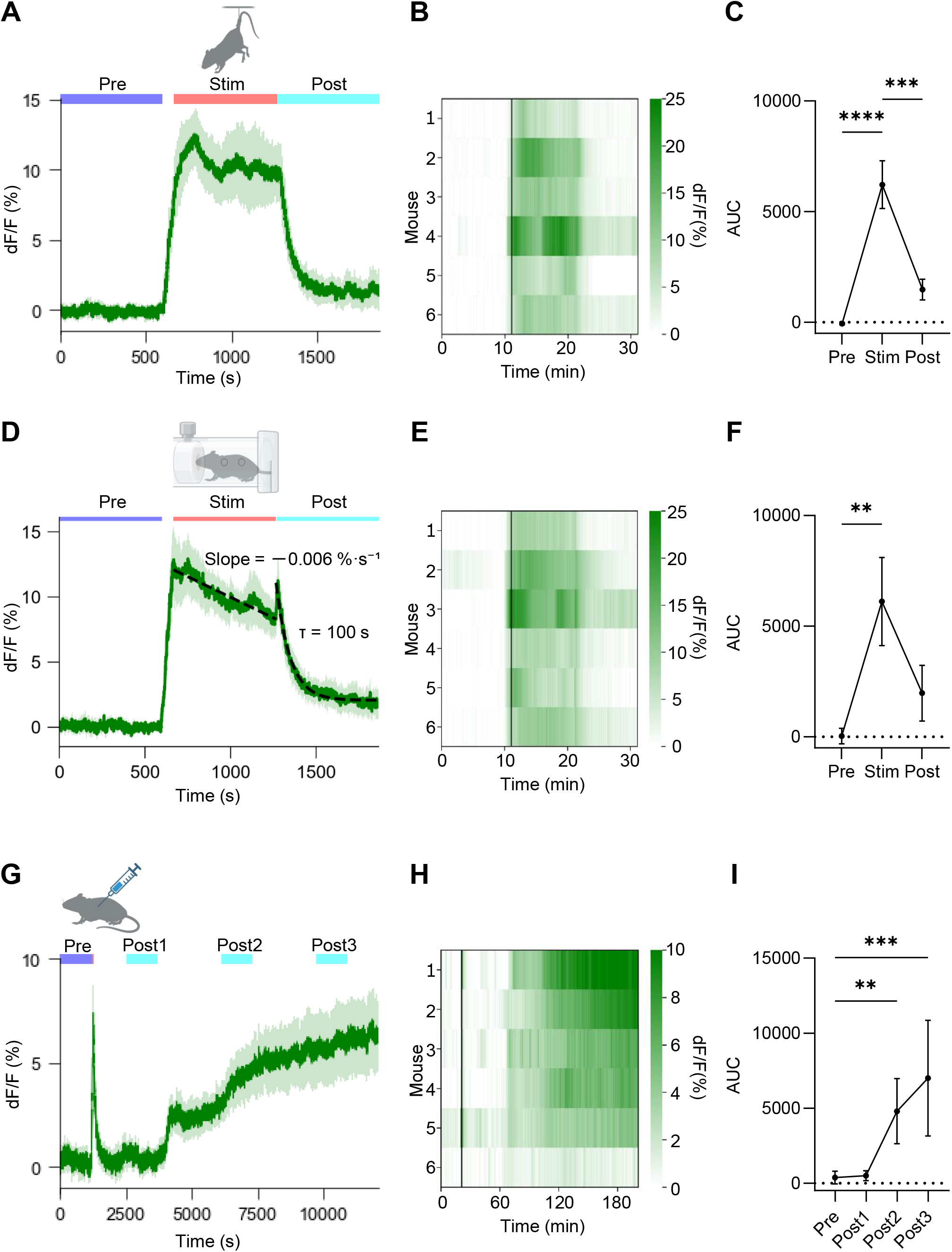
Sustained stress induces NA release on minute-to-hour timescales. A. Averaged trace showing NA release pattern during tail suspension. (Male, *N* = 3; female, *N* = 3; mean ± SEM) B. Heat map showing individual NA release during tail suspension. C. AUC values for the Pre, Stim, and Post time bins. (Male, *N* = 3; female, *N* = 3; mean ± SEM; one-way ANOVA; \*\*\**P* < 0.001, \*\*\*\**P* < 0.0001) D. Averaged trace showing NA release pattern during restraint stress. (Male, *N* = 4; female, *N* = 2; mean ± SEM) E. Heat map showing individual NA release during restraint stress. F. AUC values for the Pre, Stim, and Post time bins. (Male, *N* = 4; female, *N* = 2; mean ± SEM; Friedman’s test; \*\**P* < 0.01) G. Averaged trace showing NA release pattern following LPS administration. (Male, *N* = 4; female, *N* = 2; mean ± SEM) H. Heat map showing individual NA release following LPS administration (*i.p.*, 0.5 mg/kg). I. AUC values for the Pre, Post1, Post2, and Post3 time bins. (Male, *N* = 4; female, *N* = 2; mean ± SEM; one-way ANOVA; \*\**P* < 0.01, \*\*\**P* < 0.001)

In the restraint stress experiment, mice were restrained in a cylindrical restrainer for 10 min (Stim period) (Figure. 3D). Similar to tail suspension, NA release progressively increased from the onset of restraint and persisted throughout the stress period; however, unlike tail suspension, the signal decreased by approximately 4% immediately following the peak (Figure 3D, E). During the restraint period, the signal exhibited a shallow linear decline (slope = −0.006 %·s⁻¹). After termination of restraint stress, NA release gradually decayed with a single-exponential time constant of 100 s (Figure 3D). AUC analysis revealed that the Stim AUC was significantly greater than the Pre AUC (Figure 3F).

We also examined the effects of inflammatory stress induced by lipopolysaccharide (LPS). LPS (0.5 mg/kg) was administered intraperitoneally following a 20-min baseline recording period, and NA release was subsequently monitored for 3 h. The initial 20-min baseline period, and NA release was subsequently monitored for 3h (Figure 3G). A transient increase in NA release lasting approximately 30 s was observed at the time of injection. Subsequently, NA release gradually increased and continued to rise throughout the 3-h recording period (Figure 3G, H). The AUCs of Post2 and Post3 were significantly greater than that of the Pre period (Figure 3I). Together, these data indicate that NA release dynamics in the PVN span distinct timescales depending on the nature of the stressor, with tail suspension and restraint stress eliciting minute-scale responses and LPS producing sustained elevation over several hours.

### Specificity of the fluorescent signal

Fluorescence signals recorded by fibre photometry can be affected by motion artifacts^24^ and autofluorescent substances present in the blood^25^. To verify signal specificity, we employed pharmacological inhibition of the GRAB_NA_ sensor. The GRAB_NA_ sensor is engineered based on an α2-adrenergic receptor, and fluorescence responses have been reported to be abolished by administration of yohimbine, a selective α2-adrenergic receptor antagonist^13^. Following this approach, mice were subjected to a 1-min tail suspension before and after yohimbine administration, and fluorescence responses were compared. Comparison of signal changes induced by tail suspension before (Pre) and 30 min after yohimbine administration (Post) confirmed that the fluorescent signal increased by approximately 8% during the Pre-condition, whereas negligible changes were observed in the Post condition (Figure 4A). AUC quantification revealed that the Post AUC was significantly reduced compared with the Pre AUC (Figure 4B).

**Figure 4:**
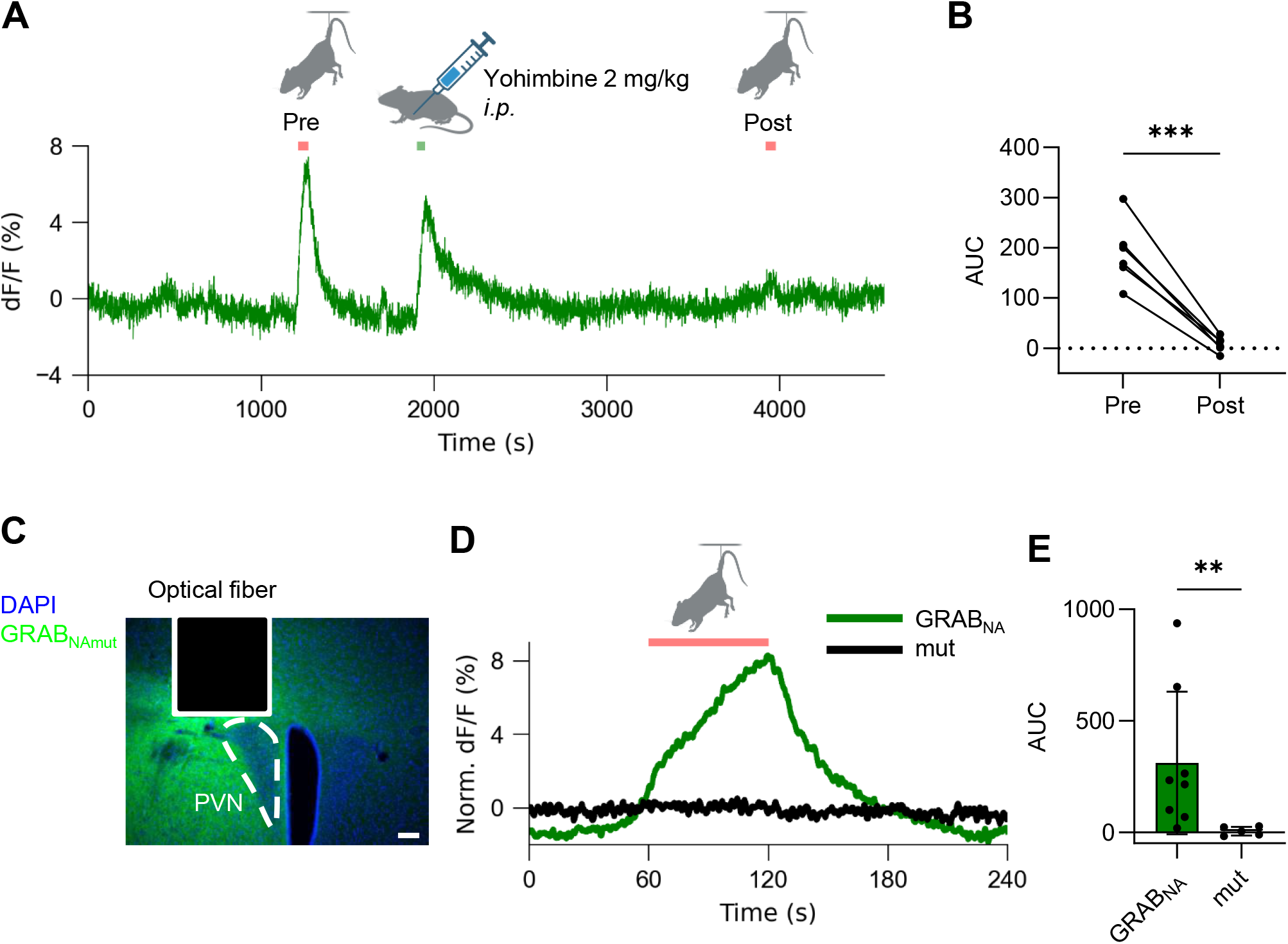
Validation of the NA specificity of signal fluctuations. A. Representative trace showing signal fluctuations during tail suspension under yohimbine treatment (*i.p.*, 2.0 mg/kg). B. AUC during tail suspension before and after yohimbine administration. (*N* = 6; paired *t*-test; \*\*\**P* < 0.001) C. Representative image showing the expression of GRAB_NAmut_ and the fibre tract. Scale bar = 100 μm. D. Averaged traces showing signal fluctuations in the GRAB_NA_ and mut groups. (GRAB_NA_, *N* = 6; mut, *N* = 5) Each trace was normalized at the onset of tail suspension (average of 59 to 60 s). E. AUC during tail suspension in the GRAB_NA_ and mut groups. (GRAB_NA_, *N* = 6; mut, *N* = 5; unpaired *t*-test; \*\**P* < 0.01)

As yohimbine is known to alter blood flow through blockade of vascular α2-adrenergic receptors, we additionally performed control experiments using the GRAB_NAmut_ sensor. GRAB_NAmut_ contains mutations in the ligand-binding domain of the GRAB_NA_ sensor that prevent NA binding. Consequently, GRAB_NAmut_ does not exhibit NA-dependent fluorescence changes whilst retaining fluorescence intensity and membrane trafficking properties comparable to those of GRAB_NA_. Mice expressing either GRAB_NA_ or GRAB_NAmut_ (Figure 4C) were subjected to a 1-min tail suspension, and fluorescence responses were compared between groups. In the GRAB_NA_ group, tail suspension induced an approximately 8% increase in fluorescence, whereas almost no fluorescence change was observed in the GRAB_NAmut_ group (Figure 4D). Consistently, AUC analysis revealed that fluorescence responses in the GRAB_NAmut_ group were significantly smaller than those in the GRAB_NA_ group during tail suspension (Figure 4E).

Taken together, these pharmacological and genetic control experiments validate the specificity of the GRABNA fluorescence signal as a reliable indicator of NA release *in vivo*.

### The release of noradrenaline in the PVN is associated with changes in animals’ behaviour

The tail suspension test is widely used as a behavioural assay for depressive-like behaviour in rodent models^26^. To examine the relationship between NA release and behavioural changes, we quantified behavioural measures, specifically struggling behaviour, during tail suspension (Figure 3A; 10 min Stim period). Periods during which mice exhibited struggling behaviour were defined as the Mobile state, whereas periods without struggling were defined as the Immobile state (Figure 5A). Comparison of the slopes of GRAB_NA_ fluorescence changes during these behavioural states revealed that fluorescence signals increased during Mobile periods and decreased during Immobile periods, indicating a close temporal relationship between NA release and behavioural state (Figure 5B). To further investigate the temporal relationship between NA release and behavioural state transitions, struggling behaviour was quantified by manually tracking animals’ nose position to estimate the distance moved. Behavioural state transitions consistently preceded the increase in NA release by approximately 0.5 s (Figure 5C). It is well established that increased immobility during the tail suspension test is indicative of stronger depressive-like behaviour^26^. We therefore analyzed the correlation between the AUC of NA release during tail suspension and immobility duration. A significant negative correlation was observed between NA release and immobility duration (Figure 5D), suggesting that increased NA release is associated with enhanced behavioural activity and may reflect a reduction in depressive-like behaviour.

**Figure 5:**
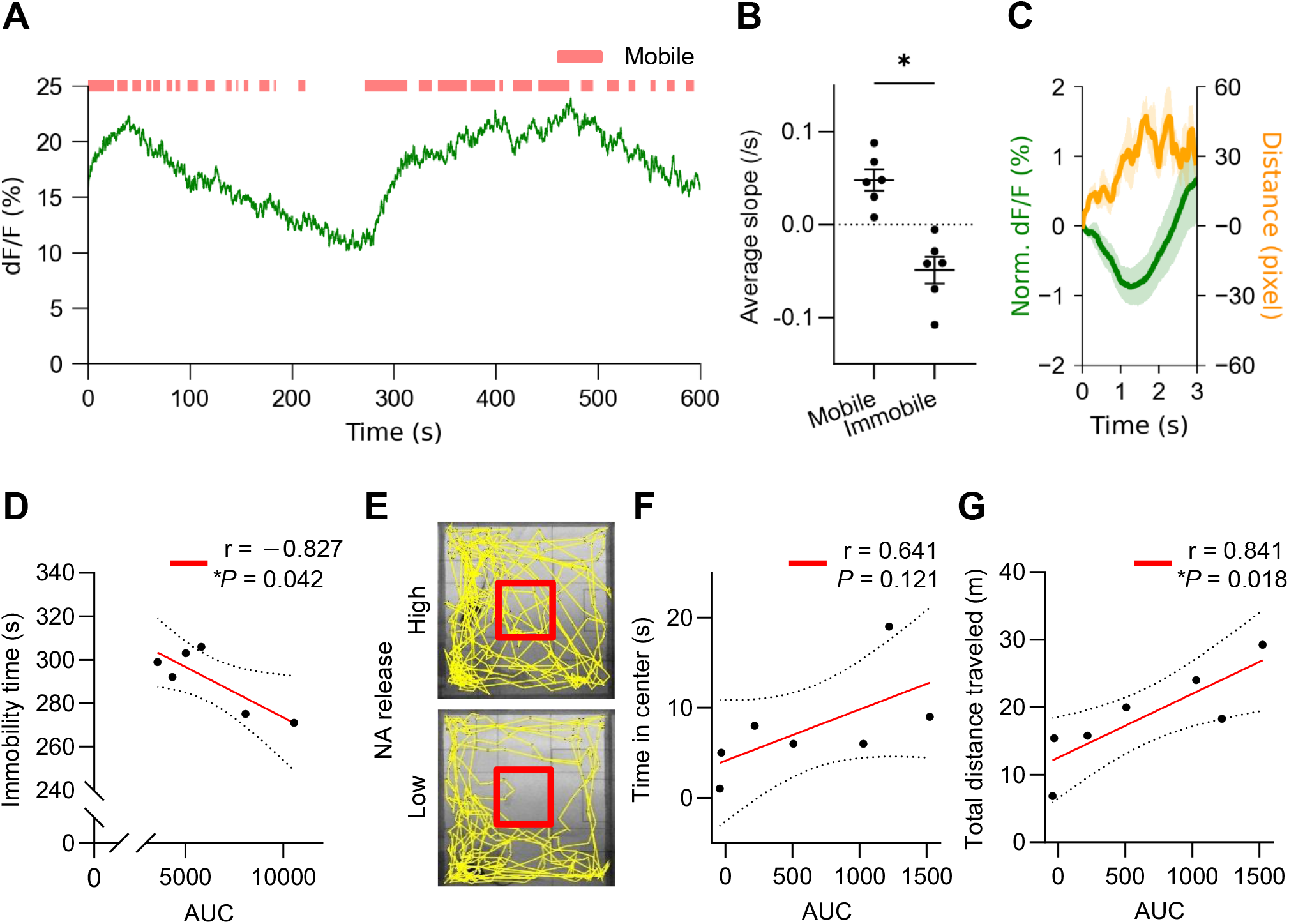
Increased NA release is associated with changes in animals’ behaviour. A. Representative trace showing NA release during tail suspension. The red bar above the trace indicates periods during which the mouse was in Mobile state. B. Weighted mean slopes of NA release during the Mobile and Immobile states. (Male, *N* = 3; female, *N* = 3; paired *t*-test; \**P* < 0.05) C. Distance travelled by the mouses’ nose (orange) and NA release (green). D. Linear regression analysis of the relationship between NA release AUC and immobility time using Pearson’s correlation coefficient. E. Representative trajectories of mice during the open-field test. F. Linear regression analysis of the relationship between NA release AUC and centre time using Pearson’s correlation coefficient. G. Linear regression analysis of the relationship between NA release AUC and total distance travelled using Pearson’s correlation coefficient.

To further investigate the relationship between NA release and anxiety-like behaviour, we performed the open-field test, a widely used assay for anxiety-related behaviour. In this test, mice with higher anxiety levels tend to avoid the centre area of the arena. We therefore analyzed the correlation between the AUC of NA release during open-field exploration and the time spent in the centre zone. A trend towards increased centre zone time with greater AUC values was observed, although this did not reach statistical significance (Figure 5E, F).

To examine the relationship between locomotor activity and NA release, we further analyzed the correlation between NA release AUC and total distance travelled during the open-field test. A significant positive correlation was identified (Figure 5G). Collectively, these findings indicate that elevated NA release in the PVN is positively associated with spontaneous locomotor activity and inversely associated with anxiety-like behaviour.

## 4. Discussion

The physiological stress response emerges from a temporally resolved regulation of both autonomic and neuroendocrine pathways^27^. In the present study, we examined the dynamics of NA release in the PVN *in vivo*, for the first time, and showed that the pattern of NA release differs, depending upon the type of stressor. Consistent with previous reports^10,28^, the majority of noradrenergic projections to the PVN originated from the NTS. Therefore, the NA release observed in the PVN in the present study is likely to be primarily derived from NTS noradrenergic neurons. Both foot shock and tail pinch, representing acute nociceptive stress, evoked rapid increases in NA release on a timescale of seconds. When animals underwent a series of foot shocks, NA release increased immediately following each stimulus delivery, whereas in the tail pinch paradigm, NA release gradually increased approximately 10 s prior to stimulus onset. The earlier rise onset in the tail pinch paradigm may be attributed to an anticipation of nociceptive stimulation and heightened alertness elicited by the approaching object^29^. Indeed, previous studies have reported that ablation of noradrenergic projections originating from the NTS suppresses psychological stress–induced increases in plasma corticosterone levels^30^. The observed rapid and synchronous NA signaling is well suited to immediately enhancing arousal and attention in response to sudden external threats, which may increase the excitability of neural circuits involved in rapid escape behaviour.

Sustained stressors such as tail suspension and restraint produced prolonged elevations in NA release that persisted throughout the duration of the stress exposure. Sustained activation of CRH neurons is thought to promote long-term increases in downstream glucocorticoid secretion, which may potentially contribute to the development of stress-related disorders. Therefore, persistently elevated NA release may preferentially act on CRH neuron populations involved not only in endocrine regulation but also in emotional processing and behavioural responses, including anxiety-, depression-, and escape-related circuits that are functionally distinct from neuroendocrine CRH neuron populations^29^.

In the tail suspension test and the open-field test, we found that increased NA release was associated with enhanced behavioural activity and negatively correlated with depression-like behaviour in mice. These findings are consistent with previous studies reporting that patients with depression exhibit lower brain noradrenaline levels than healthy individuals^31^, suggesting that reduced activity of noradrenergic projections from the NTS to the PVN contributes to the expression of depressive-like behavior. In contrast, the open-field test revealed a negative correlation between NA release and anxiety-like behaviour. This finding suggests that anxiety-like behaviour in mice is regulated not only by NA but also by other monoaminergic systems. Besides NA, serotonin has been strongly implicated in anxiety disorders^32^. Serotonin has been reported to activate CRH neurons via 5-HT2 and 5-HT7 receptors expressed in CRH neurons^13,33^. Furthermore, individuals with higher anxiety levels have been reported to exhibit greater serotonin release in the PVN^34^, suggesting that serotonin may promote anxiety through activation of CRH signaling pathways.

Intraperitoneal administration of LPS induced a gradual increase in NA release commencing approximately 15 min after injection, which continued to increase in a stepwise manner over several hours. Notably, increases in NA release were consistently observed across animals at approximately 15, 50, and 80 min after administration. This stepwise increase is distinct from the rapid, pulsatile responses observed following exposure to acute nociceptive stressors and sustained elevations during physical restraint, highlighting the unique temporal signature of inflammatory stress. Sustained NA release under inflammatory conditions may preferentially drive prolonged activation of CRH neurons, potentially contributing to the chronic dysregulation of the HPA axis observed in inflammatory diseases and stress-related psychiatric disorders.

Prostaglandin E2 (PGE2), a lipid mediator, has been reported to increase in the hypothalamus following NA administration^35^. PGE2 not only regulates hypothalamic neuronal activity^36^ but is also known to increase progressively during inflammatory responses^37^. In addition, the inflammatory cytokine interleukin-1β (IL-1β) has been shown to activate PVN neuronal activity approximately 1 h later via pathways involving the vagus nerve and NTS^38^. Therefore, the prolonged increase in NA release likely reflects the integrated actions of multiple inflammatory mediators acting across distinct timescales. Furthermore, intraperitoneal LPS administration has been reported to elevate body temperature over approximately 24 h, and this effect is attenuated by the cyclooxygenase (COX) inhibitor indomethacin^39^, suggesting that secondary prostaglandin production and ongoing inflammatory progression persist over a prolonged timescale, potentially driving sustained NA release. Future studies employing COX inhibitors, interleukin receptor antagonists, or vagotomy will be necessary to directly determine the contribution of specific inflammatory mediators at each time point and to elucidate the molecular mechanisms underlying the stepwise increase in NA release.

Together, these data reveal the unique temporal characteristics of NA release within the PVN in response to diverse stressors. The stressor-specific NA release patterns were associated with distinct behavioural outputs and anxiety levels. Future studies directly associating these dynamic NA release patterns with ensuing CRH neuron activity and downstream neuroendocrine outputs, such as corticosterone secretion, will be warranted to fully elucidate the functional consequences of stressor-specific NA signaling. This dynamic regulation of CRH neuron activity in the PVN is predicted to promote precise temporal tuning of the physiological stress response, thereby facilitating adaptive responses to environmental challenges.

## Acknowledgements

The authors thank Mr. Samuel Mestern for proofreading and editorial assistance with the manuscript. The authors are grateful to Ms. Hideko Fukuda, Ms. Masayo Igarashi, and Mr. Yoshihisa Kaneko for their dedicated support in maintaining the animal colony used in this study.

Funding: This work was supported by grants from the Japan Science and Technology Agency (JST) (JPMJCR21P1; JPMJMS2292) to T. Sasaki; the Japan Society for Promotion of Science (JSPS) (24K11683) to K. Itoi; JSPS (24K18360), the Konica Minolta Imaging Science Encouragement Award from Konica Minolta Science and Technology Foundation, the Suzuken Memorial Foundation grant, and the Yamaguchi Educational and Scholarship Foundation to H. Igarashi.

## Author contributions

Conceptualization: R.O., T.S., and H.I. Methodology: R.O., K.I., K.S., T.S., and H.I. Investigation: R.O. and H.I. Resources: K.I., K.S., T.S., and H.I. Data curation: R.O., K.I., T.S., and H.I. Validation: R.O. and H.I. Formal analysis: R.O. and H.I. Project administration: K.I., T.S., and H.I. Visualization: R.O. and H.I. Funding acquisition: K.I., T.S., and H.I. Supervision: K.I., T.S., and H.I. Writing—original draft: R.O. and H.I. Writing—review and editing: R.O., K.I., K.S., T.S., and H.I.

## Notes

### Competing Interest Statement

The authors have declared no competing interest.

