## Supplemental Figures for "Stressor-specific dynamic patterns of noradrenaline release in the paraventricular nucleus of the hypothalamus in freely moving mice"

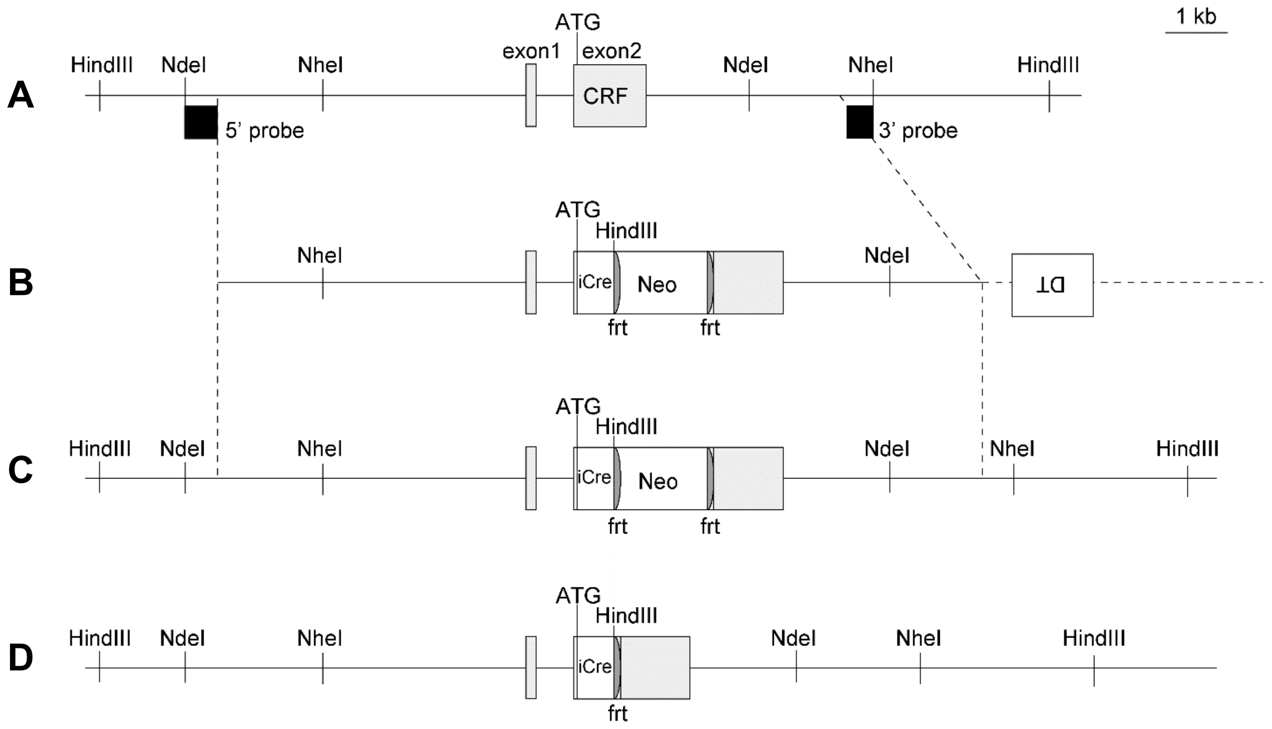


### Supplementary Figure 1: Generation of CRF-iCreΔNeo mouse

The CRF-iCre mouse carries the iCre[16] gene in the CRF genomic locus of the C57BL/6N mouse[17,18]. The targeted CRF gene locus of the CRF-iCre construct is shown in Supplementary figure 1A. The targeting vector used for homologous recombination contained exons 1 and 2, as well as the intron, of the CRF gene, and the iCre gene was inserted at the translation start site of the CRF gene in exon 2. The Neo cassette is flanked by FLP recognition target (frt) sites and located downstream of the iCre gene (Supplementary figures 1B, C). A heterozygous CRF-iCre mouse was crossed with an FLP deleter mouse (Actb-FLPe mouse)[19] to yield the CRF-iCreΔNeo mouse in which the Neo cassette was deleted from the genome of the CRF-iCre mouse (Supplementary figure 1D). The CRF-iCreΔNeo mouse was already utilized in a previous manuscript[20]. All experiments were performed according to the guidelines of the animal welfare committees of Tohoku University and Niigata University.
